# Structural characterisation of a spontaneous site 1 opening event in the insulin receptor

**DOI:** 10.64898/2026.05.31.729040

**Authors:** Amanda D. Stange, Anna Duncan

## Abstract

The insulin receptor samples multiple conformational states during ligand binding and activation, but the transient structural transitions connecting experimentally resolved receptor conformations remain poorly characterised. Here, we report a spontaneous opening event of insulin receptor site 1 observed during an unbiased molecular dynamics simulation initiated from the experimentally resolved singly insulin-bound IR1 receptor structure. The transition was characterised by separation of the L1 and FnIII-2 domains, rearrangement of site 1 contacts, and bending of the αCT segment on the initially unoccupied receptor protomer. Structural comparison of the resulting conformation revealed similarity to the asymmetric IR2-A1 and IR2-A2 receptor states associated with hybrid insulin binding sites. Together, these findings suggest that conformations compatible with hybrid-site receptor states can emerge spontaneously from the intrinsic dynamics of the insulin receptor ectodomain, supporting a conformational selection model for receptor ligand engagement.

## INTRODUCTION

The insulin receptor (IR) is a homodimeric receptor tyrosine kinase that regulates metabolic and mitogenic signalling through insulin-dependent conformational rearrangements [1], [2], [3], [4]. Structural studies over the last decade have revealed multiple ligand-bound conformations of the receptor ectodomain, providing substantial insight into the structural basis of receptor activation and ligand recognition [5], [6], [7], [8]. However, these experimentally resolved structures primarily represent stable conformational states and provide limited information about the transient structural fluctuations connecting them.

Insulin binding to IR is mediated primarily through the canonical site 1 interface formed by the L1 domain and the αCT segment [9], [10], [11]. Site 1 constitutes the high-affinity insulin binding site and plays a central role in receptor activation Recent cryo-electron microscopy studies have identified several asymmetric and symmetric insulin receptor conformations associated with different ligand occupancies. In the singly insulin-bound (IR1) structure (PDB: 7STI), one protomer adopts a characteristic site 1-bound conformation whereas the opposing protomer remains in a more *apo*-like state [7], [12]. By contrast, the IR2-S structure contains insulin bound symmetrically at both canonical site 1 interfaces, while the asymmetric IR2-A1 (PDB: 7STK) and IR2-A2 (PDB: 7STJ) conformations contain an insulin molecule in the canonical site and a second insulin molecule occupying hybrid binding sites formed through contributions from both the canonical site 1 and site 2 interfaces[7].

Despite these structural advances, the conformational transitions connecting these experimentally resolved receptor states remain poorly understood. In several receptor conformations, accessibility of the site 1 interface is influenced by surrounding domain packing and inter-domain interactions [7], [12], suggesting that local conformational rearrangements may be required to permit transitions between *apo*-like, site 1-bound, and hybrid-site receptor states. However, the extent to which these conformations can be sampled spontaneously in the absence of ligand binding at the hybrid site remains unclear.

Molecular dynamics (MD) simulations provide a means of characterising transient conformational fluctuations and heterogeneous conformational ensembles that are difficult to resolve experimentally[13], [14]. However, large-scale spontaneous transitions remain challenging to capture in unbiased simulations because such events are often rare and occur on long timescales[14], [15], [16]. Nevertheless, rare conformational transitions observed in unbiased trajectories can provide insight into intrinsic structural tendencies and metastable conformations accessible to the receptor[17], [18].

Here, we report a spontaneous opening event of insulin receptor site 1 observed during an unbiased MD simulation initiated from the experimentally resolved singly insulin-bound IR1 receptor structure. The transition was characterised by progressive separation of the L1 and FnIII-2 domains, rearrangement of the αCT-associated interface, and localised disruption of contacts surrounding site 1. Structural comparison of the resulting conformation revealed similarity to the asymmetric IR2-A1 and IR2-A2 receptor states, suggesting that conformations associated with hybrid-site receptor structures are intrinsically accessible within the conformational landscape of the insulin receptor ectodomain.

## RESULTS

### Observation of spontaneous opening of insulin receptor site 1

To investigate the conformational stability of insulin receptor site 1, we analysed a set of unbiased molecular dynamics simulations initiated from a singly insulin-bound receptor conformation. During one of the simulations, one receptor protomer underwent a spontaneous transition associated with progressive separation of the L1 and FnIII-2 domains and rearrangement of the site 1 interface (Figure 1 and Figure S1).

**Figure 1.**
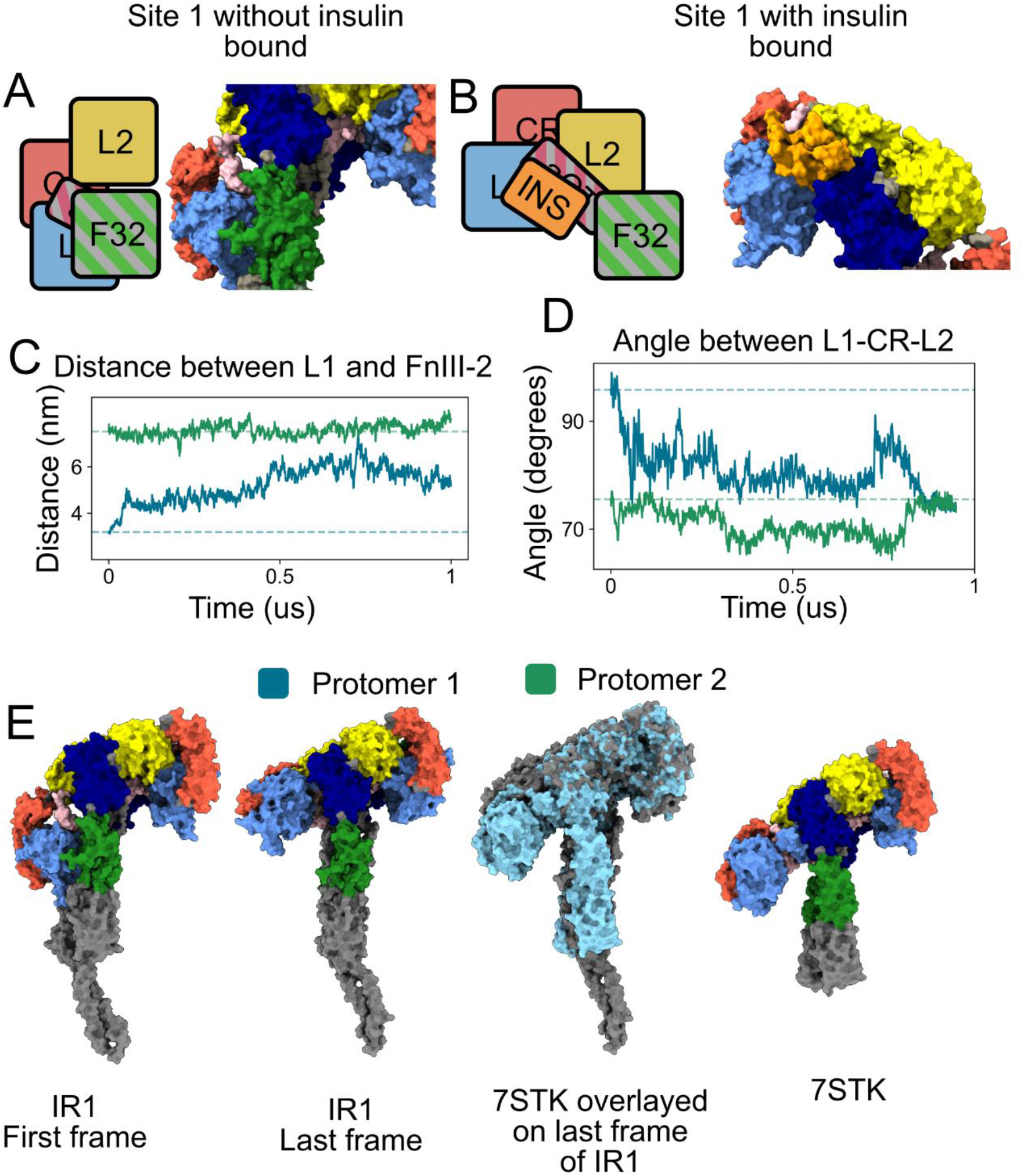
Spontaneous opening of insulin receptor site 1 during unbiased molecular dynamics simulation. (A) Schematic and structural representation of the site 1 interface in the absence of insulin binding in IR1. The L1 domain is in pale blue, L2 in yellow, CR in orange. The FnIII-2 domain of the opposing protomer is shown in green (and with grey stripes in the schematic), the FnIII-1 domain in dark blue, and the opposing αCT domain is shown in pink (with grey stripes in the schematic). (B) Schematic and structural representation of site 1 with insulin bound in IR1. Insulin is shown in orange, and other domains are coloured as in (A). (C) Distance between the COM of L1 and FnIII-2 domains during the simulation for each receptor protomer. Dashed lines indicate the distances in the initial structure. The distances for all repeats are shown in Figure S1B. (D) Angle formed between the L1, CR, and L2 domains throughout the simulation for each protomer. Dashed lines indicate the corresponding angles in the initial structure. The angles for all repeats are shown in Figure S1C. (E) Representative receptor conformations showing the first and last simulation frames together with structural comparison to the asymmetric insulin receptor conformation IR2-A1 (PDB: 7STK)[7]. The overlay demonstrates similarity between the opened simulation conformation and the experimentally resolved asymmetric receptor state. Domains are coloured as in (A). The RMSD of the two protomers for all 4 repeats are shown in Figure S1A.

For much of the simulation, both receptor protomers remained in conformations resembling the initial closed state. However, one protomer transitioned toward a more open conformation characterised by displacement of the L1 domain away from the FnIII-2 region (Figures 1C and 1D). The transition emerged after extended sampling of the closed state and occurred without externally applied biasing forces.

The opening transition was accompanied by increasing separation between the L1 and FnIII-2 domains together with changes in the relative orientation of the L1, CR, and L2 domains (Figures 1C and 1D). By contrast, the second protomer remained comparatively stable throughout the trajectory and largely retained conformations similar to the starting structure (Figure S2).

Structural comparison of the final simulation frame with experimentally resolved insulin receptor conformations revealed similarity to the asymmetric IR2-A1 structure (PDB: 7STK), particularly in the relative positioning of the L1 and FnIII-2 regions (Figure 1E). The transition therefore represents spontaneous sampling of a partially opened site 1 conformation within an unbiased simulation trajectory.

### Contact characterisation of the conformational transition

To characterise the structural rearrangements associated with spontaneous opening of site 1, we analysed residue-residue contacts that were disrupted or formed during the transition (Figure 2). Contact occupancies were compared before and after the opening event together with occupancies from the remaining simulation repeats, in which no spontaneous opening of site 1 occured.

**Figure 2.**
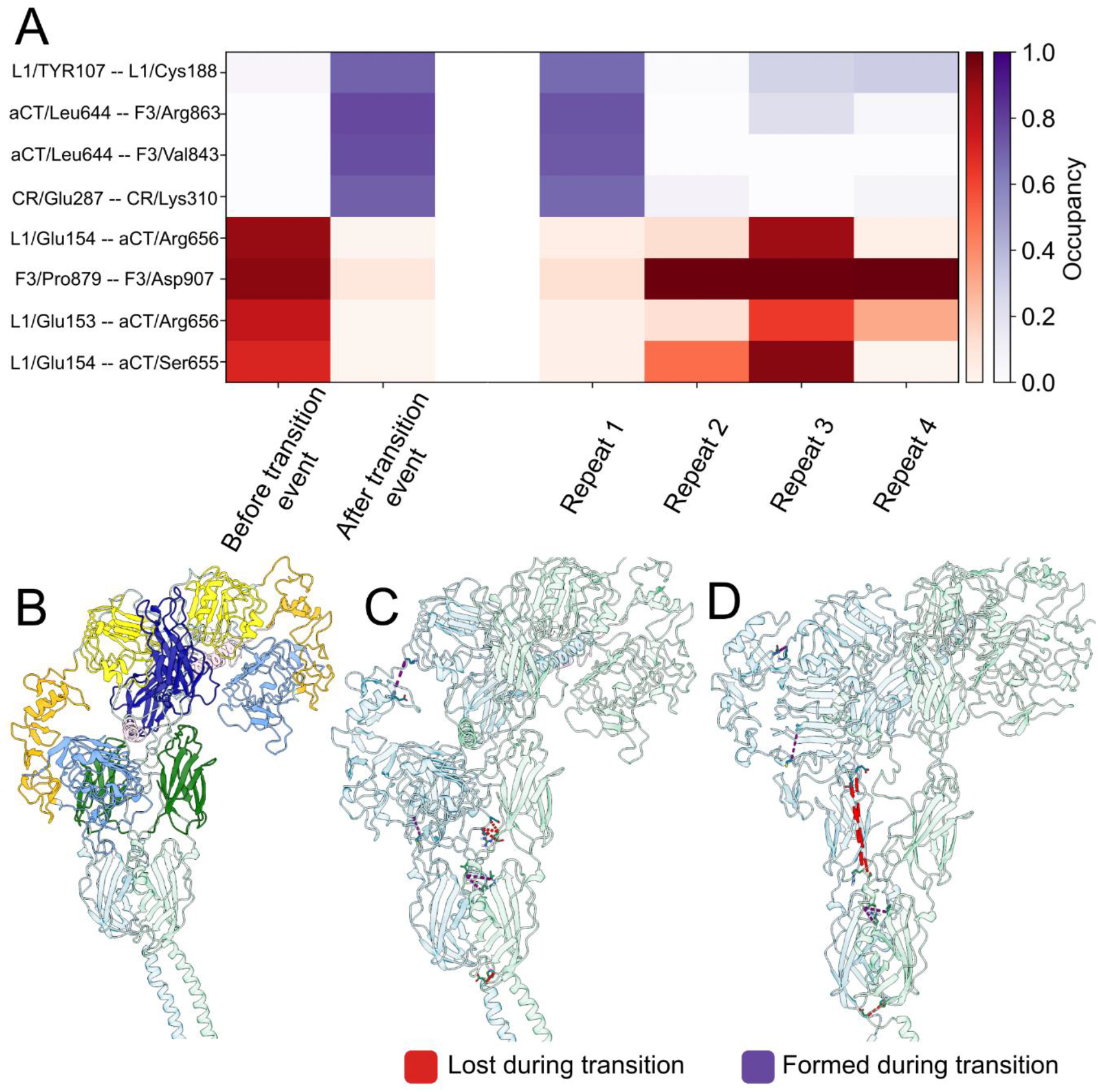
Contact rearrangements associated with spontaneous opening of insulin receptor site 1. (A) Contact occupancies for selected residue-residue interactions associated with the opening transition. Contacts are grouped according to interactions lost during the transition (red), and contacts formed during the transition (purple). Occupancies are shown before and after the transition together (which occurred in Repeat 1) alongside occupancies from Repeat 1 (over the entirety of the simulation) and the remaining simulation repeats. The selected contacts are shown as a function of time for all 4 repeats in Figure S2. (D) Initial structure of the receptor coloured by domains. The first protomer is shown in pale blue and the second protomer in pale green. The L1 domain is in pale blue, L2 in yellow, CR in orange. The FnIII-2 domain of the opposing protomer is shown in green (and with grey stripes in the schematic), the FnIII-1 domain in dark blue, and the opposing αCT domain is shown in pink. (C)Structural mapping of contacts associated with the initial stages of the opening transition. Lost contacts are shown in red, and newly formed contacts in purple. The first protomer is shown in pale blue and the second protomer in pale green. (D)Structural mapping of contacts following progression toward the more open conformation. Lost contacts are shown in red, and newly formed contacts in purple. The protein is coloured as in (B).

The opening transition was associated with disruption of multiple contacts linking the L1 domain to neighboring regions surrounding the site 1 interface (Figure 2A). Several interactions displayed high occupancy prior to the transition but were substantially weakened or absent following opening. These disrupted contacts were localised primarily to interfaces involving the L1 domain, FnIII-2 region, and neighboring structural elements.

In addition to contact loss, the transition trajectory sampled a small number of interactions that were absent or only weakly populated in the remaining simulations (Figure 2A). Several new contacts also formed during progression toward the opened conformation, indicating local reorganisation of the surrounding interface rather than complete destabilisation of the receptor structure.

Mapping the contact rearrangements onto the receptor structure showed that the largest structural changes remained localised primarily to the site 1 region and adjacent inter-domain interfaces (Figures 2C and 2D).

#### αCT rearrangements

The spontaneous opening transition was accompanied by conformational rearrangement of the αCT segment (Figure 3). Prior to the transition, both αCT segments remained predominantly extended, with bending angles centered near ∼160–170° (Figure 3A). During the transition, one protomer underwent a substantial decrease in αCT angle, corresponding to progressive bending of the helix, while the second protomer (with insulin bound) remained comparatively stable throughout the simulation.

**Figure 3.**
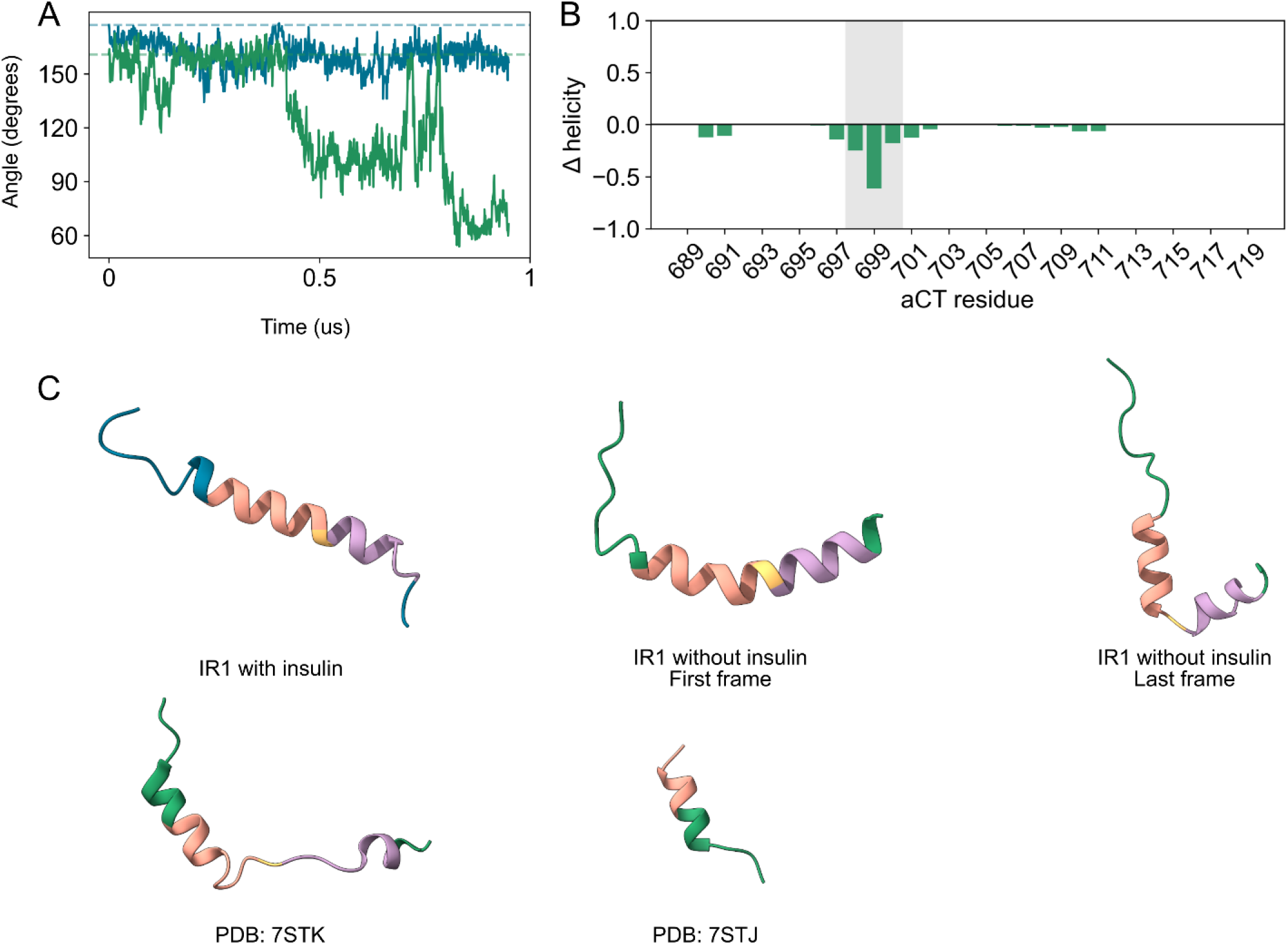
αCT rearrangement during spontaneous opening of insulin receptor site 1. (A)Time evolution of the αCT bending angle in the opening and non-opening receptor protomers. (B) Change in residue-wise αCT helicity following the opening transition relative to the initial state. (C) Representative conformations of the αCT segment before and after opening compared with experimentally resolved receptor conformations (PDB: 7STK and 7STJ [7]).

Residue-wise secondary structure analysis showed localised loss of helicity primarily within the central region of the αCT segment, centered around residues 697–700 (Figure 3B). The largest reduction in helicity was observed near residue 699, whereas the remainder of the helix largely retained its secondary structure.

Representative structures from the trajectory demonstrated progressive bending of the αCT segment during the opening event (Figure 3C). The final simulation frame adopted a conformation resembling experimentally resolved insulin receptor structures including IR2-A1 (PDB: 7STK), while remaining less bent than the αCT conformation observed in IR2-A2 (PDB: 7STJ)[7].

## DISCUSSION

In this study, we observed a spontaneous opening event of insulin receptor site 1 during an unbiased molecular dynamics simulation initiated from the experimentally resolved singly insulin-bound receptor structure IR1 (PDB: 7STI). The transition was characterised by progressive separation of the L1 and FnIII-2 domains, rearrangement of contacts surrounding site 1, and conformational bending of the αCT segment. Structural comparison of the resulting conformation revealed similarity to the asymmetric IR2-A1 and IR2-A2 receptor states, indicating that conformations associated with hybrid-site receptor structures are intrinsically accessible within the conformational landscape of the insulin receptor ectodomain.

Recent structural studies have substantially expanded the number of experimentally resolved insulin receptor conformations and highlighted the structural plasticity of the receptor ectodomain. However, these structures primarily represent relatively stable conformational states and provide limited information about the transient structural fluctuations connecting them. The spontaneous opening event observed here demonstrates that substantial rearrangements of the site 1 interface can occur directly within an unbiased simulation trajectory without externally applied forces or enhanced sampling approaches.

Experimentally resolved insulin receptor structures containing insulin bound at the canonical site 1 interface display a broadly conserved protomer architecture despite differences in overall ligand occupancy and receptor asymmetry. In the IR1 structure, one protomer adopts this characteristic site 1-bound conformation, whereas the opposing protomer, which lacks bound insulin, remains in a more *apo*-like state. During the present simulation, the site 1-bound protomer remained comparatively stable throughout the trajectory and retained a conformation similar to the initial experimentally resolved structure. By contrast, the unoccupied protomer underwent progressive opening characterised by displacement of the L1 domain, rearrangement of site 1 contacts, and bending of the αCT segment.

Structural comparison of the final simulation frame revealed that the rearranged protomer adopts a conformation resembling the asymmetric IR2-A1 and IR2-A2 receptor states, particularly in the relative positioning of the L1 and FnIII-2 regions and the accompanying αCT rearrangement. In these experimentally resolved structures, insulin occupies hybrid binding sites formed through contributions from both the canonical site 1 and site 2 interfaces. The spontaneous transition observed here therefore shifts the initially apo-like protomer of IR1 towards a conformation compatible with experimentally observed hybrid-site receptor states.

The observed transition is consistent with a conformational selection model for insulin receptor dynamics, in which the receptor intrinsically samples conformational states associated with later stages of ligand engagement prior to insulin binding at the hybrid site (Figure 4). In this framework, insulin binding would preferentially stabilise conformations already accessible within the intrinsic receptor ensemble rather than inducing the conformational change entirely through ligand engagement. The spontaneous emergence of an IR2-A1/A2-like conformation in the absence of insulin bound at the hybrid site therefore suggests that these asymmetric receptor states may arise from pre-existing conformational tendencies within the ectodomain.

**Figure 4.**
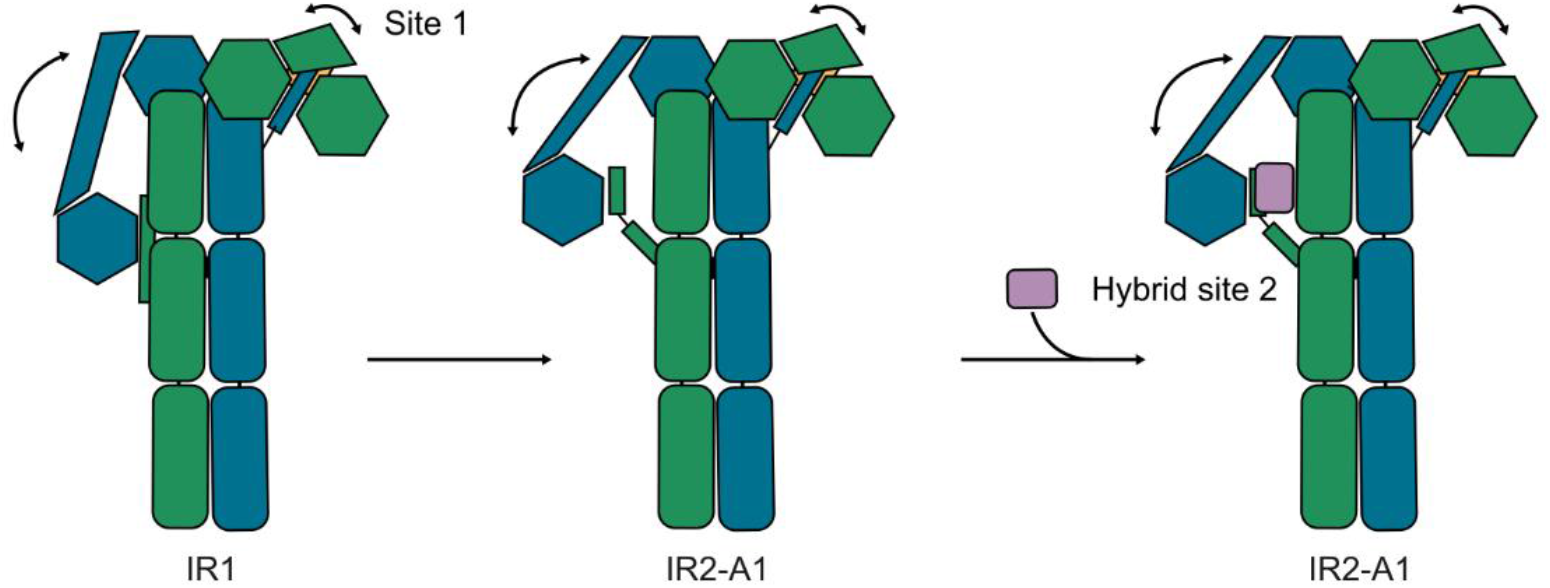
Hypothesis of opening of site 1 before insulin binds.

The αCT rearrangements observed during the transition further support the dynamic nature of the site 1 interface. Opening was accompanied by localised loss of helicity and progressive bending within the central region of the αCT segment, while much of the remaining helix remained preserved. These conformational changes resemble features observed in experimentally resolved asymmetric receptor conformations and suggest that αCT flexibility contributes to transitions between apo-like and hybrid-site-compatible receptor states.

Together, these findings demonstrate that the insulin receptor ectodomain can spontaneously sample conformations associated with experimentally resolved hybrid-site receptor states within an unbiased molecular dynamics trajectory and provide a structural description of the contact rearrangements and αCT conformational changes underlying this rare transition.

### Limitations of the study

The spontaneous opening event described here was observed in a single unbiased molecular dynamics trajectory and therefore does not permit quantitative assessment of the frequency, kinetics, or thermodynamic stability of the observed conformational transition. Although the transition occurred without externally applied biasing forces, the stochastic nature of rare conformational events makes it difficult to determine how frequently similar rearrangements occur within the broader receptor conformational ensemble.

As with all molecular dynamics studies, the observed behaviour may depend on the selected force field, membrane composition, simulation conditions, and starting structure. The simulations were initiated from a specific singly insulin-bound receptor conformation, which may itself preferentially stabilise particular regions of conformational space.

The present work focuses specifically on structural rearrangements associated with spontaneous opening of the site 1 interface and does not directly address the functional consequences of the observed transition for receptor activation, ligand exchange, negative cooperativity, or downstream signalling.

Furthermore, while the opened conformation exhibited similarity to experimentally resolved asymmetric receptor states, the physiological relevance of the observed transition remains to be established experimentally.

Additional computational and experimental studies will therefore be required to determine the prevalence and functional significance of transient site 1 opening events within the insulin receptor conformational landscape.

## RESOURCE AVAILABILITY

### Lead contact

Requests for further information and resources should be directed to and will be fulfilled by the lead contact, Anna Duncan.

### Materials availability

This study did not generate new unique reagents.

### Data and code availability

Simulation coordinate data have been deposited at Zenodo and are publicly available as of the date of publication, the DOI is listed in the key resources table. Any additional information required to reanalyze the data reported in this paper is available from the lead contact upon request.

## ACKNOWLEDGMENTS

We had access to computational resources at the Grendel cluster of the Centre for Scientific Computing Aarhus, and the Resource for Biomolecular Simulations (ROBUST; supported by the Novo Nordisk Foundation; NNF24OC0087976).

## AUTHOR CONTRIBUTIONS

Conceptualisation, A.D.S. and A.D.; methodology, A.D.S.; investigation, A.D.S.; formal analysis, A.D.S.; molecular dynamics simulations, A.D.S.; writing – original draft, A.D.S.; writing – review & editing, A.D.S.,and A.D.; supervision, A.D.; funding acquisition, A.D.

## DECLARATION OF GENERATIVE AI AND AI-ASSISTED TECHNOLOGIES IN THE WRITING PROCESS

During the preparation of this work, the authors used ChatGPT (OpenAI) to assist with language editing and manuscript structuring. All scientific content, analysis, interpretation, and final editing were performed and verified by the authors.

## SUPPLEMENTAL INFORMATION

**Figure S1.**
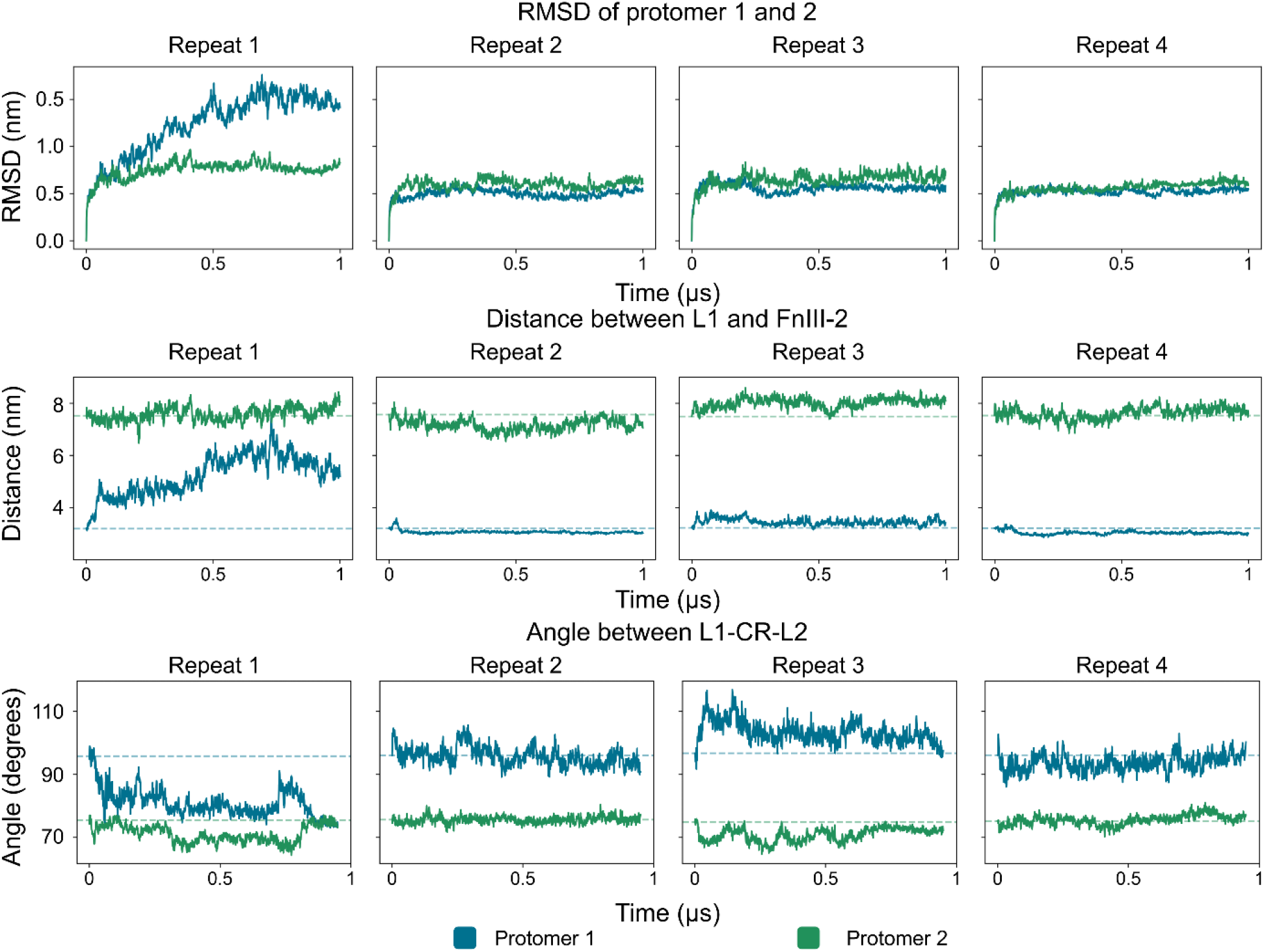
RMSD and L1 progression for all repeats. (A) The RMSD of protomer 1 and 2 as a function of time for the 4 repeats. Protomer is coloured in blue and protomer 2 in green. (B) The distance between the COM of the L1 and FnIII-2 domains as a function of time for the 4 repeats. Protomer is coloured in blue and protomer 2 in green. (C) The angle between the COMs of the L1, CR, and L2 domains as a function of time for the 4 repeats. Protomer is coloured in blue and protomer 2 in green.

**Figure S2.**
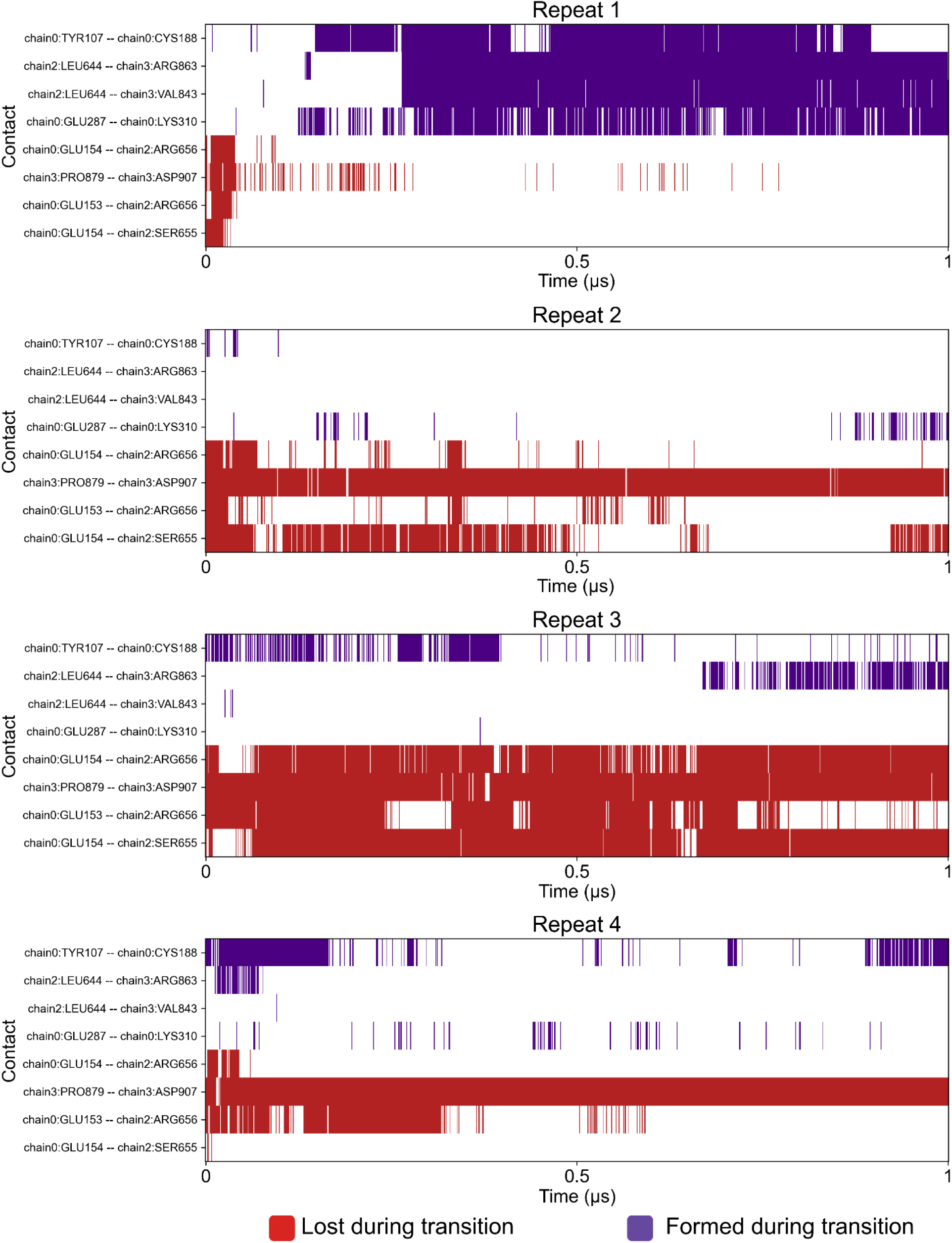
Selected contacts as a function of time. Contacts lost during the transition is coloured in red and formed coloured in purple.

## STAR★METHODS

### KEY RESOURCES TABLE

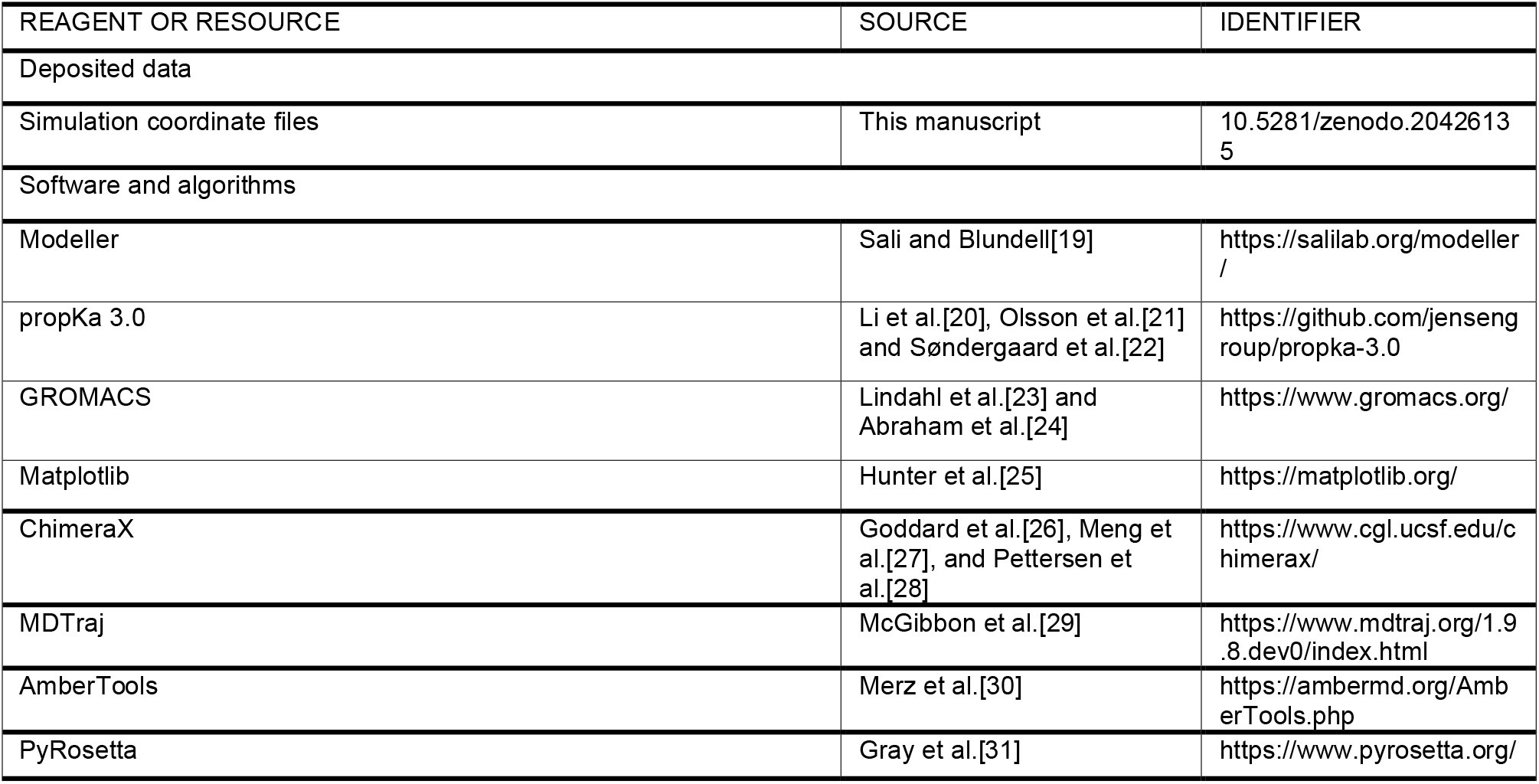

## METHOD DETAILS

### Molecular dynamics simulations

Initial full-length insulin receptor models were generated using Boltz2[32] based on the experimentally resolved IR1 ectodomain structure (PDB 7STI). Predicted models were refined using Modeller[19] and the relax protocol[33], [34], [35], [36] in PyRosetta[31], [37] prior to MD simulations. Transmembrane (TM) dimer conformations were clustered based on TM helix geometry, and representative models from the dominant interface states were selected for simulation.

Simulation systems were prepared using the AABY pipeline (available at AABY GitHub), which integrates COBY for membrane construction and protein insertion[38], tleap for generation of AMBER-format force field parameters[30], ParmEd for conversion of AMBER topologies to GROMACS-compatible formats[39], and GROMACS tools for solvation and ion placement[23], [24]. The receptor was embedded in a lipid bilayer composed of 60% POPC and 40% cholesterol, solvated using OPC water, and neutralised with 0.15 M NaCl. Protein parameters were assigned using the AMBER19SB force field and lipid parameters using Lipid21[40], [41].

Energy minimisation was followed by multi-stage equilibration with gradually released positional restraints. Production MD simulations were performed in GROMACS using a 2 fs timestep. Four independent 1 μs simulations were performed. Simulations were carried out at 310 K using the velocity-rescale thermostat and at 1 bar using the C-rescale barostat with semi-isotropic pressure coupling[42], [43]. Long-range electrostatics were treated using particle mesh Ewald, with 0.9 nm real-space electrostatic and van der Waals cut-offs. Bonds involving hydrogen atoms were constrained using the LINCS algorithm[44]. Coordinates were saved every 1 ns for analysis.

For the analyses presented in this study, trajectories corresponding to TM interface cluster 2 were analysed in detail due to the presence of spontaneous Site 1 opening transitions.

### Trajectory analysis

Trajectory analysis was performed using MDTraj[29], GROMACS tools[23], and custom Python scripts. Structural rearrangements associated with Site 1 opening were characterised using domain centre-of-mass (COM) distances, inter-domain angles, residue contact analyses, and αCT conformational metrics.

Domain COMs were calculated using MDTraj (md.compute_center_of_mass) from backbone atoms within the L1 (residues 1–192), CR (195–310), L2 (313–446), FnIII-1 (474–590), FnIII-2 (596–637 and 754–772), and FnIII-3 (776–875) domains. Inter-domain angles were derived from vectors connecting domain COMs to quantify large-scale ectodomain rearrangements during opening transitions.

Residue contacts were calculated using MDTraj minimum heavy-atom distance metrics (md.compute_contacts, scheme = closest-heavy). Contacts were represented as binary occupancy matrices using a 0.45 nm distance threshold. For the rare-event trajectory, pre- and post-transition occupancies were calculated relative to frame 50. This cutoff was chosen as it represents the initial jump in the L1-FnIII-2 distance as seen in Figure 1C. Lost contacts were defined as contacts with pre-transition occupancy ≥0.40 and post-transition occupancy ≤0.20, whereas formed contacts were defined as contacts with pre-transition occupancy ≤0.20 and post-transition occupancy ≥0.40. This initially led to 14 contacts from where 8 was found to have a higher occupancy in the post-transition frames compared to the pre-transition and 5 found to have a higher occupancy in the pre-transition frames. From this selection 4 of the post-transition contacts were removed as they had an occupancy higher than 0.40 in at least one of the 3 repeats that did not undergo a transition, and one of the pre-transition contacts was removed as this had an occupancy lower than 0.40 in all the repeats that did not undergo a transition. This left a total of 8 contacts that were identified to be different pre-and post-transition and between repeat 1 and repeat 2-4.

αCT helix bending was quantified using a kink angle calculated between two fitted helical axes. For each trajectory frame, principal axes were fitted by singular value decomposition to backbone atoms from the N-terminal αCT segment (residues 691–698) and the C-terminal αCT segment (700–709). The N-terminal axis was oriented from residue 698 to 691 and the C-terminal axis from residue 700 to 709 to ensure consistent vector directionality. The kink angle was defined as the angle between the two segment axes for each αCT chain.

αCT secondary structure was analysed using DSSP as implemented in MDTraj (md.compute_dssp). The two αCT helices were analysed separately using residues 689–719 from the corresponding receptor chains. DSSP assignments were converted into binary helix occupancies by assigning residues classified as α-helical (H) a value of 1 and all other secondary-structure states a value of 0. Per-residue helix occupancies were calculated over the first and last 100 ns of each trajectory. Changes in helicity were reported as Δ helicity, defined as the difference between post-window and pre-window helix occupancies.

Plots were generated using matplotlib[25], and structural visualisations were produced using ChimeraX[26], [27], [28].

## QUANTIFICATION AND STATISTICAL ANALYSIS

Trajectory analysis and data processing were performed using custom Python scripts together with MDTraj[29], NumPy[45], pandas[46], [47], SciPy[48], and matplotlib[25]. Structural visualisation was performed using ChimeraX[26], [27], [28].

For molecular dynamics simulations, *n* refers to independent simulation replicas unless otherwise stated. Four independent replicas were simulated. Replicas were prepared independently with distinct initial lipid, solvent, and ion configurations.

No formal statistical hypothesis testing was performed in this study. All analyses were based on descriptive comparisons of independently simulated trajectories.

